# Toward Large-Scale Preclinical Neuroimaging: Quantifying Inter-Scanner Variability in Mouse Brain MRI

**DOI:** 10.64898/2026.05.29.728711

**Authors:** Moussa Shahid, Jiangyang Zhang

**Affiliations:** Department of Chemistry, Hunter College, New York, NY, USA; Center for Biomedical Imaging, Department of Radiology, New York University Grossman School of Medicine, New York, NY, USA

## Abstract

Multi-site MRI studies in preclinical neuroimaging are emerging, but unlike in human studies, characterization of inter-scanner variability remains limited. In this study, we assessed intra- and inter-scanner variability between two similarly equipped 7 Tesla MRI scanners using a phantom and ex vivo mouse brain specimens. Diffusion-weighted imaging revealed slight differences in gradient amplitudes between the scanners, while estimated apparent diffusion coefficient (ADC) values showed a coefficient of variation below 1.5% and inter-scanner differences below 2% near the magnet center. Volumetric analysis based on proton density-weighted images showed negligible intra-scanner differences across sessions, while inter-scanner volumetric differences were mostly less than 2% and spatially non-uniform across the brain. Quantitative maps of R1, R2*, and MTsat showed inter-scanner relative differences of less than 5%, 10%, and 20%, respectively, with white matter exhibiting greater variability than gray matter. These findings provide a foundation for future large-scale, multi-scanner preclinical neuroimaging studies.

## 1. Introduction

Magnetic resonance imaging (MRI) is a non-invasive neuroimaging tool used both clinically and in the study of laboratory animal models of neurological disorders, including multiple sclerosis, stroke, and Alzheimer’s disease. In these models, MRI facilitates the investigation of disease mechanisms and the evaluation of potential treatments. Its suitability for brain imaging stems from several key advantages: unlike conventional optical and ultrasound imaging, MRI is unimpeded by the skull, and unlike X-ray and computed tomography, it offers a broad range of soft tissue contrasts with greater sensitivity to subtle changes in tissue microstructure. Compared to conventional histology, which yields rich cellular detail but is often limited in scope due to the time-consuming processes of sectioning and staining, MRI enables whole-brain survey and produces data that are readily quantified. Although the development of modern 3D light sheet microscopy after tissue clearing has alleviated some of these limitations of conventional histology, challenges remain for many protocols, including lengthy sample preparation, incomplete antibody penetration, and tissue deformation depending on the clearing method used. In this context, MRI can provide fast, non-destructive macroscopic characterization of the whole brain, guiding targeted follow-up with conventional histology or 3D light sheet microscopy for cellular-level detail, and thereby improving overall throughput for characterizing disease progression in animal models.

Most studies using MRI to examine animal models have been conducted by a single group on a dedicated MRI scanner. This potentially limits the throughput for large scale studies involving comparisons across multiple models or cohorts, and multi-site studies in this domain are only beginning to emerge. By contrast, pooling data across multiple sites and large cohorts is standard practice in human brain imaging. Large-scale multi-site initiatives, such as the Human Connectome Project (HCP) and the UK Biobank, have demonstrated that aggregating neuroimaging data increases statistical power and enables the detection of subtle disease-related brain changes that would otherwise remain undetectable in single-site studies ([1]; [2]). As the field moves toward large-scale, multi-site collaborations, correcting for inter-site differences becomes essential ([3]). These site-specific differences arise from multiple sources: variations in scanner hardware such as gradient coil design, RF coil geometry, and main field homogeneity; and differences in acquisition sequences and parameters; and reconstruction algorithms. Left uncorrected, these effects hinder large-scale multi-site studies by artificially inflating variance within and across groups, reducing sensitivity to detect real biological differences, and introducing site or scanner-related confounds ([4]). These problems have been studied extensively in human MRI data, where researchers employ both prospective and retrospective strategies to mitigate their impact ([5]), whereas efforts to address them in animal MRI remain limited, largely due to the lack of multi-site studies before.

In human neuroimaging, several strategies have been developed to address this problem. For example, in prospective protocol harmonization, where sites agree on identical acquisition parameters across vendors before any data are collected ([6]). Also human studies often use Traveling Human Phantoms, where the same group of volunteers is scanned at every site to quantify scanner bias ([7]). A third approach is retrospective statistical harmonization, in which algorithms such as ComBat are applied after acquisition to remove site or scanner-related variance while preserving meaningful biological signals ([8]). Initiatives such as ADNI have combined these strategies to ensure data comparability across major vendors ([9]).

In preclinical MRI, retrospective harmonization is generally not feasible, as most sites lack sufficient data acquired under comparable protocols. In this study, we characterized inter-scanner differences between two similarly equipped 7 Tesla (T) MRI scanners using a phantom and ex vivo mouse brain specimens. We employed a quantitative MRI protocol sensitive to both macroscopic brain morphology, assessed through volumetric measures, and tissue microstructure. These findings represent a first step toward the design of future large-scale, multi-scanner cohort studies in preclinical neuroimaging.

## 2. Materials and Methods

### 2.1 Experimental setup, phantom, and ex vivo mouse brain specimens

All animal experiments have been approved by the Institute Animal Care and Use Committee at New York University. Adult C57BL/6 mice (7-9 weeks old, n = 3, 2M/1F, Jackson Laboratory, Bar Harbor, ME, USA) were perfusion fixed with 4% paraformaldehyde in PBS. The samples were preserved in 4% PFA for 24 h before transferring to PBS. The specimens were imaged with the skull intact and placed in a syringe filled with Fomblin (perfluorinated polyether, Solvay Specialty Polymers USA, LLC, Alpharetta, GA, USA) to prevent tissue dehydration. To test the performance of the gradient systems, a phantom was created by filling a 10 mL syringe with silicon coil, which has lower diffusivity than water at room temperature.

Experiments were performed on two similarly equipped 7 T horizontal MRI scanners (Bruker Biospin, Billerica, MA, USA). Scanner 1 was equipped with a gradient system capable of generating a maximum gradient strength of 660 mT/m, and Scanner 2 was equipped with a gradient system capable of generating a maximum gradient strength of 550 mT/m. Both scanners used an 82-mm diameter circularly polarized birdcage radiofrequency resonator (Bruker Biospin, Ettlingen, Germany) for homogeneous signal transmission, in conjunction with a four-channel receive-only phased array CryoProbe (CRP) anatomically shaped to closely conform to the rodent head (Bruker Biospin, Ettlingen, Germany) for high receiver sensitivity. The CRP used with Scanner 1 was designed for mouse brain imaging, while the one used with scanner 2 was designed for rat brain imaging and was slightly larger than the CRP on scanner 1.

### 2.2 MRI Data Acquisition

MRI data were acquired using a quantitative MRI protocol. Proton density (PD)-weighted, T1-weighted, and magnetization transfer (MT)-weighted images were acquired using a 3D multiple gradient echo sequence with the following parameters: first echo time (TE)/repetition time (TR) = 2.5 ms/50 ms, six echoes with echo spacing of 2.5 ms, flip angle of 6◦, 21◦, and 6◦, respectively, and a spatial resolution of 0.15 mm x 0.15 mm x 0.15 mm, 2 signal average. For the MT-weighted image, a 2kHz off-resonant radio frequency pulse was used with a pulse duration of 4 ms and an amplitude of 10 uT. The total imaging time to acquire all three images was approximately 1 hour. The experiment was repeated 5 times on each scanner to measure intra-scanner variability.

### 2.3 MRI data analysis

Proton density (PD)-, T1-, and MT-weighted images were analyzed using the QUIT toolbox (https://github.com/spinicist/quit) to generate quantitative R1, R2*, and MTsat maps. To compare parameter maps across sessions and scanners, all images were registered to a common template using affine and nonlinear registrations implemented in ANTs. Global and voxel-wise Jacobian maps were then derived from these transformations. The Jacobian quantifies local volumetric changes when one image is deformed to match another: a value of 1 indicates no change in local volume, values greater than 1 indicate expansion, and values between 0 and 1 indicate compression. Taking the natural logarithm of the Jacobian yields the log-Jacobian, which re-centers the scale at zero, such that a value of 0 indicates no volume change, positive values indicate expansion, and negative values indicate compression. This logarithmic transformation makes the metric symmetric around zero, facilitating a more intuitive interpretation of volumetric differences than the raw Jacobian.

### 2.4 Statistical Analysis

To examine whether there was significant differences between measurements from the two scanners, we performed pair t-test with the threshold for statistical significance set at 0.05.

## 3. Results

### 3.1 Comparisons of imaging gradient performance

Diffusion-weighted signals from the phantom revealed slight differences in gradient amplitudes between the two scanners (**Fig. 1A**). However, the estimated ADC values showed no significant difference between them (**Fig. 1B**). Along the direction of the main magnetic field (B_0_), inter-scanner differences in estimated ADC remained insignificant, though they increased slightly with distance from the center. Overall, the coefficient of variation in ADC was less than 1.5% within 8 mm of the magnet center, and the inter-scanner difference in ADC was less than 2%.

**Figure 1:**
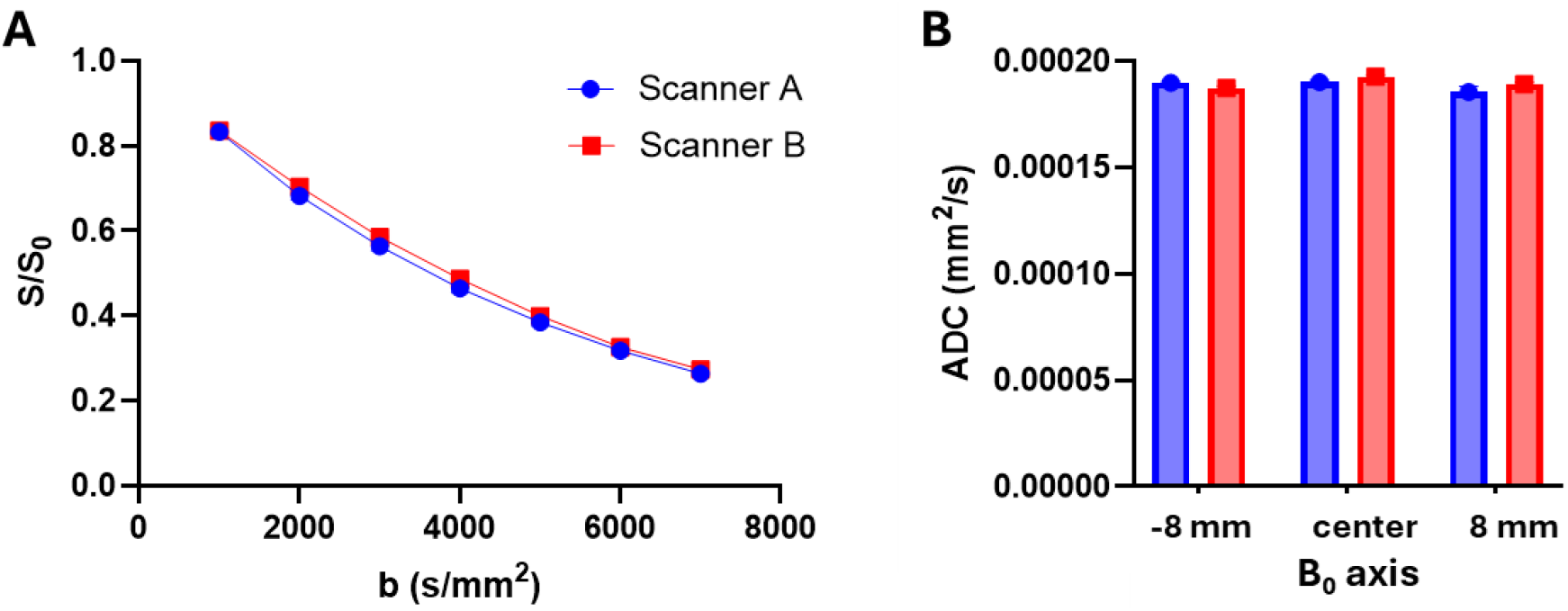
Comparison of gradient system performance between the two scanners. (A) Average diffusion-weighted signal attenuation curves measured at the isocenter of both scanners. (B) Estimated ADC values at three locations along the z-direction (B_0_) for both scanners. In both panels, data points represent the means and error bars indicate the standard deviations of five repeated measurements.

### 3.2 Inter-scanner differences in mouse brain morphology

Based on proton density-weighted images collected from the same mouse brains (**Fig. 2**), volumetric changes between images acquired on the same scanner across different sessions were negligible. In contrast, volumetric changes between images acquired on the two different scanners were present, though mostly less than 2%. The log-Jacobian map (**Fig. 2**) reveals that these volumetric changes were spatially non-uniform across the brain. In this representative brain, the dorsal region showed slight enlargement, while larger local volumetric changes were more prevalent in regions outside the brain.

**Figure 2:**
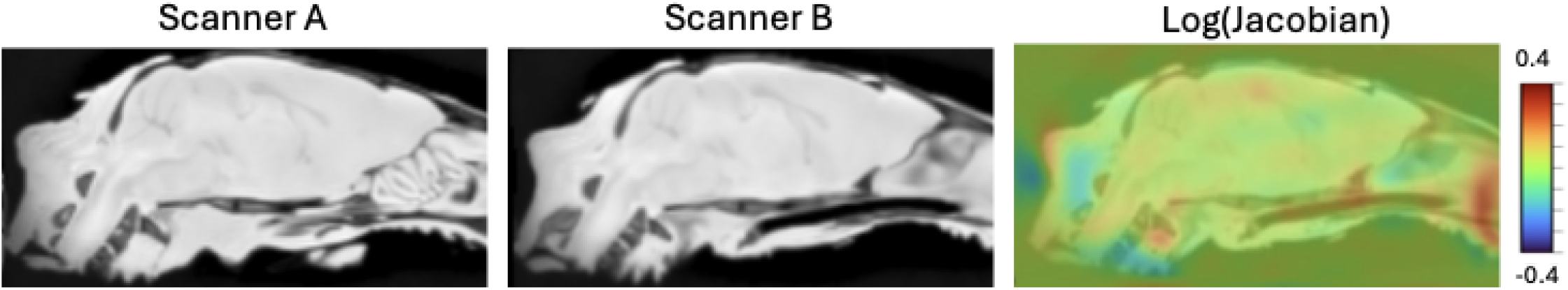
Representative proton density weighted images from the same mouse brain scanned on both scanners and the voxel-wise Jacobian map showing local volumetric changes between the two images.

### 3.3 Inter-scanner differences in mouse brain tissue MR properties

Quantitative maps of R1, R2*, and MTsat acquired on both scanners were compared to assess intra- and inter-scanner differences (Fig. 3). One scanner performed more consistently, with discrepancies ranging from approximately 2–4% across all three maps, indicating greater intra-scanner stability. The other scanner exhibited both higher and more variable discrepancies, with intra-scanner relative differences in R1 and R2* ranging from 1–4% and MTsat reaching as high as 9% — roughly two to four times the values observed with the first scanner. As expected, inter-scanner differences were noticeably larger than intra-scanner differences, with estimated relative differences of less than 5% for R1, less than 10% for R2*, and less than 20% for MTsat. For all three parameter maps, both intra- and inter-scanner differences were spatially non-uniform across the brain, with white matter tending to exhibit higher differences than gray matter.

**Figure 3:**
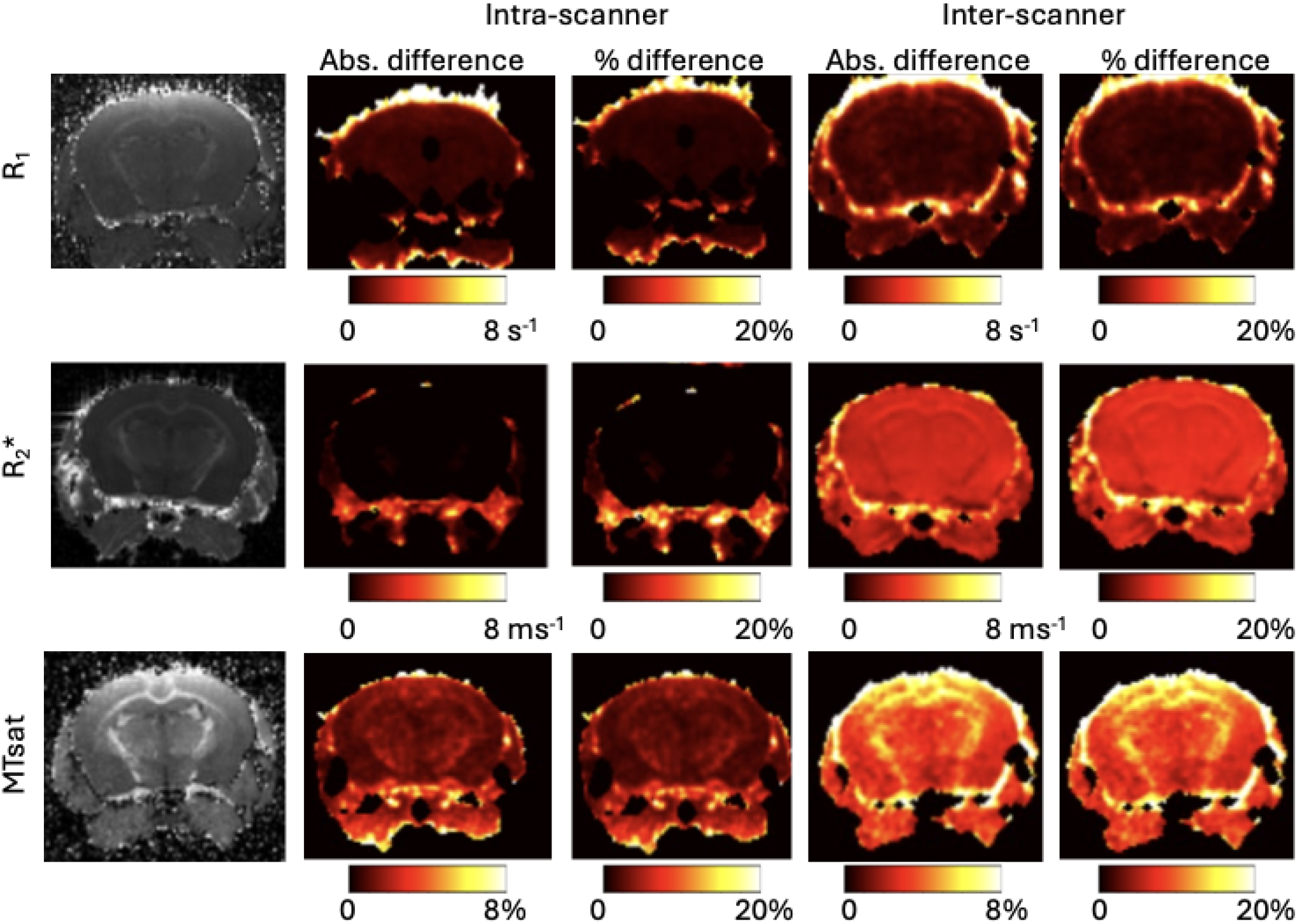
Intra-scanner and inter-scanner differences in tissue parameter maps. For both intra-scanner and inter-scanner differences, both absolute and relative differences are shown.

## Discussion

Despite the development of guidelines and tools for multi-site harmonization in human neuroimaging, equivalent resources for preclinical animal studies remain lacking. Preclinical rodent brain imaging is predominantly conducted on high-field MRI systems (7 T or higher)[10], primarily because the rodent brain is approximately 1,000 times smaller than the human brain and therefore demands higher spatial resolution. Higher field strengths enable the smaller voxel sizes required for mouse neuroanatomy, but also give rise to field-dependent artifacts, including B0 and B1 inhomogeneities, that must be carefully characterized and corrected. Furthermore, intrinsic tissue MR properties, such as T1 and T2 relaxation times, vary with field strength, adding another layer of complexity to cross-scanner harmonization [11]. In institutes operating multiple MRI scanners, the systems often differ in field strength to accommodate a broad range of applications such as spectroscopy and tumor imaging, which introduces significant technical challenges for cross-scanner comparisons.

In this study, we focused on characterizing differences between two 7 T scanners with similar hardware configurations under near-ideal conditions. To minimize the effects of B0 and B1 inhomogeneities, large-volume excitation coils and high-order shimming were employed on both scanners. Comparable receiver coils were used to minimize differences in coil geometry and sensitivity profiles. Furthermore, imaging protocols were based on the same pulse sequences with identical acquisition parameters across both scanners. Together, these measures represent a well-controlled experimental setup that isolates residual inter-scanner differences from other potential confounds.

The modest discrepancies observed in R1 and R2* can be attributed to the relatively straightforward nature of their fitting procedures. Our intra-scanner comparison revealed that R2* exhibited smaller variability than R1, likely because R2* maps were derived from six echo images, whereas R1 maps were estimated from only two images acquired at different flip angles. Conversely, R2* showed higher inter-scanner variability than R1, which may reflect differences in gradient performance or B0 inhomogeneities between the two scanners. MTsat, by contrast, exhibited larger intra- and inter-scanner variability than both R1 and R2*. MTsat variability reached as high as 9–16%, while R1 and R2* remained comparatively stable at 4–7%. This elevated variability is likely attributable to the fact that MTsat is a derived quantity that depends in part on the estimated R1, meaning that errors in R1 estimation may propagate into the MTsat estimate. This is further supported by the observation that white matter structures showed higher MTsat variability than gray matter regions. Such tissue-dependent inter-scanner variability poses additional challenges for cross-scanner data harmonization.

This study has several limitations. First, ex vivo mouse brains were used, and the resulting tissue properties may not fully reflect in vivo conditions, as postmortem changes, chemical fixation, and temperature differences can alter relaxation times, diffusion characteristics, and other MR signal properties. Second, our experimental setup represented a near-ideal scenario, with both scanners operating at the same field strength and equipped with comparable hardware; as such, the observed inter-scanner differences may underestimate the variability expected in real-world multi-site studies where scanner configurations, field strengths, and coil designs differ more substantially. Finally, we did not apply or evaluate statistical harmonization approaches, such as post hoc correction methods, that could help mitigate site-related effects while preserving biologically meaningful signals. Together, these limitations highlight important methodological considerations for improving reproducibility and comparability in future multi-site preclinical MRI studies.

In summary, we characterized intra- and inter-scanner variability in ex vivo mouse brain MRI at 7 T using a standardized quantitative MRI protocol. We found volumetric measurements between the two scanners to be highly consistent. Our results also demonstrate that while intra-scanner variability was relatively low, considerable inter-scanner variability was observed in quantitative R1, R2*, and MTsat maps, with MTsat exhibiting the largest discrepancies. These findings highlight the importance of careful scanner characterization and suggest that inter-scanner differences pose meaningful challenges for large-scale, multi-site preclinical MRI studies. Addressing these challenges will require the development of dedicated harmonization strategies tailored to the specific demands of preclinical neuroimaging.

## References

[1] Van Essen DC, Smith SM, Barch DM, Behrens TE, Yacoub E, Ugurbil K; WU-Minn HCP Consortium. The WU-Minn Human Connectome Project: an overview. Neuroimage. 2013 Oct 15;80:62–79. doi: 10.1016/j.neuroimage.2013.05.041. Epub 2013 May 16. PMID: 23684880; PMCID: PMC3724347.

[2] Miller KL, Alfaro-Almagro F, Bangerter NK, Thomas DL, Yacoub E, Xu J, Bartsch AJ, Jbabdi S, Sotiropoulos SN, Andersson JL, Griffanti L, Douaud G, Okell TW, Weale P, Dragonu I, Garratt S, Hudson S, Collins R, Jenkinson M, Matthews PM, Smith SM. Multimodal population brain imaging in the UK Biobank prospective epidemiological study. Nat Neurosci. 2016 Nov;19(11):1523–1536. doi: 10.1038/nn.4393. Epub 2016 Sep 19. PMID: 27643430; PMCID: PMC5086094.

[3] Kislik G, Fox R, Korotcov A, Zhou J, Febo M, Moghadas B, Bibic A, Zou Y, Wan J, Koehler RC, Adebayo T, Burns MP, McCabe JT, Wang KKW, Huie JR, Ferguson AR, Paydar A, Wanner IB, Harris NG and The TOP-NT Investigators (2025) Multi-site, in vivo MRI dataset of brain diffusivity measures before and after harmonization, and atrophy measures following controlled cortical impact in male and female adult rats. Front. Neurol. 16:1719618. doi: 10.3389/fneur.2025.1719618

[4] Elkabes S, Li H. Proteomic strategies in multiple sclerosis and its animal models. Proteomics Clin Appl. 2007 Oct 16;1(11):1393–1405. doi: 10.1002/prca.200700315. PMID: 19759847; PMCID: PMC2744133.

[5] Jack CR Jr, Bernstein MA, Fox NC, Thompson P, Alexander G, Harvey D, Borowski B, Britson PJ L Whitwell J, Ward C, Dale AM, Felmlee JP, Gunter JL, Hill DL, Killiany R, Schuff N, Fox-Bosetti S, Lin C, Studholme C, DeCarli CS, Krueger G, Ward HA, Metzger GJ, Scott KT, Mallozzi R, Blezek D, Levy J, Debbins JP, Fleisher AS, Albert M, Green R, Bartzokis G, Glover G, Mugler J, Weiner MW. The Alzheimer’s Disease Neuroimaging Initiative (ADNI): MRI methods. J Magn Reson Imaging. 2008 Apr;27(4):685–91. doi: 10.1002/jmri.21049. PMID: 18302232; PMCID: PMC2544629.

[6] Kim H, Lee K, Chang S, Kang G, Tak Y, Lee M, Kim V, Lee J, Jeong H. Factors affecting the validity of self-reported data on health services from the community health survey in Korea. Yonsei Med J. 2013 Jul;54(4):1040–8. doi: 10.3349/ymj.2013.54.4.1040. PMID: 23709443; PMCID: PMC3663212.

[7] Treit S, Stolz E, Rickard JN, McCreary CR, Bagshawe M, Frayne R, Lebel C, Emery D, Beaulieu C. Lifespan Volume Trajectories From Non-harmonized T1-Weighted MRI Do Not Differ After Site Correction Based on Traveling Human Phantoms. Front Neurol. 2022 May 9;13:826564. doi: 10.3389/fneur.2022.826564. PMID: 35614930; PMCID: PMC9124864.

[8] Fortin JP, Parker D, Tunç B, Watanabe T, Elliott MA, Ruparel K, Roalf DR, Satterthwaite TD, Gur RC, Gur RE, Schultz RT, Verma R, Shinohara RT. Harmonization of multi-site diffusion tensor imaging data. Neuroimage. 2017 Nov 1;161:149–170. doi: 10.1016/j.neuroimage.2017.08.047. Epub 2017 Aug 18. PMID: 28826946; PMCID: PMC5736019.

[9] Demerath EW, Sun SS, Rogers N, Lee M, Reed D, Choh AC, Couch W, Czerwinski SA, Chumlea WC, Siervogel RM, Towne B. Anatomical patterning of visceral adipose tissue: race, sex, and age variation. Obesity (Silver Spring). 2007 Dec;15(12):2984–93. doi: 10.1038/oby.2007.356. PMID: 18198307; PMCID: PMC2883307.

[10] Kreitz S, Mennecke A, Konerth L, Rösch J, Nagel AM, Laun FB, Uder M, Dörfler A, Hess A. 3T vs. 7T fMRI: capturing early human memory consolidation after motor task utilizing the observed higher functional specificity of 7T. Front Neurosci. 2023 Aug 10;17:1215400. doi: 10.3389/fnins.2023.1215400. PMID: 37638321; PMCID: PMC10448826.

[11] Spincemaille, Pascal & Anderson, Julie &Wu, Gaohong & Yang, Baolian & Fung, Maggie & Li, K. & Li, Shaojun & Gupta, Ajay & Kelley, Douglas & Benhamo, Nissim & Wang, Yi. (2019). Quantitative Susceptibility Mapping: MRI at 7T versus 3T. Journal of Neuroimaging. 30. 10.1111/jon.12669.

